# ONeSAMP 3.0: Effective Population Size via SNP Data for One Population Sample

**DOI:** 10.1101/2023.09.14.557784

**Authors:** Aaron Hong, Rebecca G. Cheek, Kingshuk Mukherjee, Isha Yooseph, Marco Oliva, Mark Heim, W. Chris Funk, David Tallmon, Christina Boucher

**Affiliations:** Department of Computer and Information Science and Engineering, University of Florida, Gainesville, FL, United States; Department of Biology, Graduate Degree Program in Ecology, Colorado State University, Fort Collins, CO, United States; Department of Math, Colorado State University, Fort Collins, CO, United States; Biology and Marine Biology Program, University of Alaska Southeast, Juneau, AK, United States

**Author notes:** These authors should be considered co-senior authors (David Tallmon, Christina Boucher).

## Abstract

1. The genetic effective size (Ne) is arguably one of the most important characteristics of a population as it impacts the rate of loss of genetic diversity. Genetic estimators of (Ne) increasingly popular tools in population and conservation genetic studies. Yet there are very few methods that can estimate the Ne from data from a single population and without extensive information about the genetics of the population, such as a linkage map, or a reference genome of the species of interest.
2. We present ONeSAMP 3.0, an algorithm for estimating Ne from single nucleotide polymorphism (SNP) data collected from a single population sample using Approximate Bayesian Computation and local linear regression.
3. We demonstrate the utility of this approach using simulated Wright-Fisher populations, and empirical data from five endangered Channel Island fox (*Urocyon littoralis*) populations to evaluate the performance of ONeSAMP 3.0 compared to a commonly used Ne estimator. Our results show that ONeSAMP 3.0 is robust to the number of individual samples and number of loci included in and appears accurate even if the range of true Ne values is large.
4. This method is broadly applicable to natural populations and is flexible enough that future versions could easily include summary statistics appropriate for a suite of biological and sampling conditions. ONeSAMP 3.0 is publicly available under the GNU license at https://github.com/AaronHong1024/ONeSAMP_3 and also available with Bioconda (https://bioconda.github.io/index.html).

## 1 Introduction

Effective population size (Ne) is one of the most important parameters in population biology because it influences a host of evolutionary processes that affect population viability [1–5]. From a broad perspective, Ne can be defined as an “ideal” population that would experience drift at the same rate as the sample population [6]. As the number of potential parents in a population decreases, or variance of individual reproductive success increases, the Ne decreases [7]. From an applied point of view, the Ne determines the rate of loss of genetic diversity due to stochastic events [8, 9], inbreeding [10], and can be impacted by demographic and biological characteristics of a species [11, 12]. Populations that have recently experienced a bottleneck event to a small Ne can have reduced fitness and elevated short-term extinction risk [13, 14]. Furthermore, Ne may also inform whether a population can maintain adequate genetic variance for adaptive evolution in response to environmental change [15, 16]. Estimating Ne is therefore of crucial importance in conservation and evolutionary biology.

The increasing availability of genetic data for many non-model organisms has catalyzed a tremendous effort in population genetics to accurately and precisely estimate Ne using molecular marker data [7]. A number of useful genetic estimators have been developed to estimate Ne (e.g. [17–19]). A few approaches for estimating current Ne have been developed to take advantage of genomic datasets from a single sample of individuals [20–22]. One highly successful one-sample Ne estimator based on gametic disequilibrium is the software `LDNe` [23], included in the software NeEstimator [24].

The use of summary statistics has been proposed as an alternative to metrics population genetics that cannot efficiently be calculated exactly [25]. These statistics offer a close approximation to their computationally-intensive counterparts, and have proved to be successful in some empirical applications [26–28]. Most applications have used a rejection sampling method [26], in which all summary statistic values that fall outside a given tolerance range are rejected, and only those summary statistics that fall within the tolerance range are used to estimate the target parameters [27]. In this paper, we present ONeSAMP 3.0, which uses local linear regression and smooth weighting to improve the accuracy and precision of parameter estimation from summary statistics within an approximate Bayesian framework [29, 30].

We combine multiple summary statistics in a Bayesian approach to capture the genotypic and allelic information contained in a single population genetic sample to estimate Ne [29, 30]. The original implementation of ONeSAMP 3.0 [30] used summary statistics and approximate Bayesian computation to estimate Ne from a single sample of microsatellite data. Here, we describe an updated and remodeled ONeSAMP 3.0 by adding new features which include the ability to accept single nucleotide polymorphism (SNP) data in GENEPOP format and data filtering algorithms to adjust for low coverage loci and individuals. We first outline the background of ONeSAMP 3.0’s approach in detail, before exploring the performance of ONeSAMP 3.0 compared to `LDNe` in replicate simulations with known Ne values and sample sizes typical of published SNP datasets. Finally, we apply both methods to an empirical dataset from Channel Island foxes to test ONeSAMP 3.0 against a dataset of sub-populations which have experienced varying levels of population bottlenecks. ONeSAMP 3.0 gives a precise and reliable estimate of Ne for recently bottlenecked populations with appropriate sampling sizes. When the number of individuals sampled is less than 100, and the Ne value is relatively large (greater than or equal to 200) then ONeSAMP 3.0 is particularly useful. ONeSAMP 3.0 was implemented in python and went through rigorous usability testing. The complied method is publicly available on Bioconda. All source code and instructions for using ONeSAMP 3.0 is available on GitHub in anticipation that it will become an important part of the toolkit for researchers studying non-model organisms with limited sample sizes and numbers of loci genotyped.

## 2 Method

### 2.1 Background

ONeSAMP 3.0 estimates the Ne of an population using a similar paradigm as ONeSAMP [30]. Briefly, it starts by simulating a large number of datasets with an identical number of individuals and loci as well as a known Ne value. Next, five statistics are computed for each of the simulated populations as well as the input population. Lastly, linear regression is used to to infer the Ne value for the input population based on the statistics of the input population, and the known Ne value and statistics for the simulated populations. We note that ONeSAMP is restricted to microsatelite data, and thus, uses a subset of the same statistics as ONeSAMP 3.0. In contrast, ONeSAMP 3.0 has been redeveloped to use SNP data in GENEPOP format. In this section, we introduce these statistics used by ONeSAMP 3.0, and then describe the Bayesian model used to estimate Ne from the simulated populations.

#### 2.1.1 Calculation of summary statistics

Any summary statistics that can be calculated from standard population genetic data could be incorporated. We limited ONeSAMP 3.0 to a few that are straightforward to calculate, commonly used in population genetics studies, and related to Ne based upon previous results and simulations of our own. The five summary statistics incorporated in the ONeSAMP 3.0 Ne estimator are calculated for the allelic or genotypic data from a sample of individuals genotyped at multiple loci. These summary statistics are defined as follows.

##### Estimated mean expected heterozygosity

The first summary statistic calculated by ONeSAMP 3.0 is the estimated mean expected heterozygosity, denoted by *Ĥ*_*e*_. It has a positive relationship with effective population size, and is the proportion of heterozygous genotypes expected under Hardy-Weinberg equilibrium [31]. *Ĥ*_*e*_ is also incorporated into many of the more complicated population genetics statistics used to infer population structure and dynamics so we begin with its definition below:

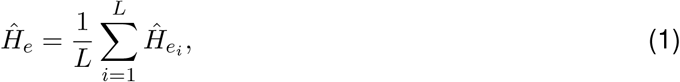

where *L* is the number of loci, and 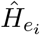 is the heterozygous frequency of allele *i* in the population.

##### Fixation index

In a very small, non-selfing population, allele frequencies can differ between males and females, causing a heterozgygote excess in their progeny over that expected based upon Hardy-Weinberg proportions. This quantity was estimated using Wright-Fisher model calculated by dividing the observed heterozygosity by the expected heterozygosity. This is also referred to as the *F*-statistic or fixation index. It is denoted as 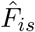 [31] and is defined as follows:

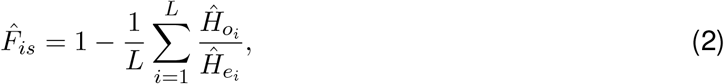

where the observed heterozygosities 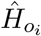 and expected heterozygosities 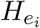 are averaged over all of the *L* loci.

#### Multi-locus homozygosity

Next, we used the first two moments of multi-locus homozygosity, the mean (denoted as *ĥ*) and variance (denoted as 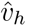), as summary statistics. These can be calculated from counting the number of loci at which each of *S* individuals in a sample are homozygous (*h*).

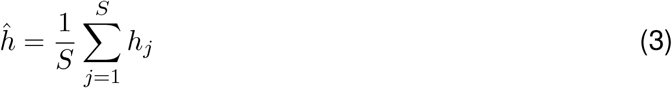

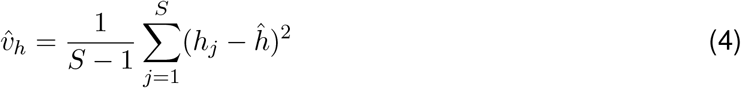

It has been shown that the moments of the distribution of an analogous measure, multi-locus heterozygosity, provide insights into Ne [32]. Higher moments of multi-locus homozygosity were not used after initial simulations suggested they were uninformative under the range of parameter values we examined.

#### Gametic disequilibrium

An estimate of gametic disequilibrium can be used to infer Ne when the reciprocal of the average of this statistic is taken over all pairs of distinct loci [33]. Non-random associations among alleles at different loci, or gametic disequilbria, are generated by finite Ne [34]. We calculate the square, 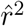, of Burrow’s estimator of gametic disequilibrium, 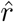, for the case where the phases of double heterozygotes are indistinguishable:

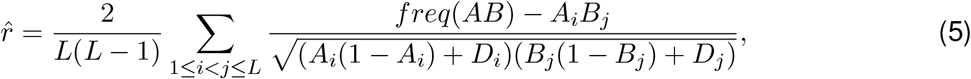

where *L* is the number of loci, *freq*(*AB*) is the frequency of locus *A* and locus *B* occurring together, *A*_*i*_ is the frequency of allele *A* at locus *i, B*_*j*_ is the frequency of allele *B* at locus *j, D*_*i*_ and *D*_*j*_ are the respective departures of the loci from Hardy-Weinberg equilibrium, and *i* = 1, …, *L* and *j* = 1, …, *L*.

#### 2.1.2 Generating simulated populations

In ONeSAMP 3.0, we make use of an individual-based Wright-Fisher simulation model of a diploid species [35] with a target Ne. This model differs slightly from a Wright-Fisher model in that there are two allogamous sexes and equal numbers of each sex. This causes Ne to slightly exceed the sum of the number of males and females in the population [36]. Given that we are trying to estimate current Ne, and the population may not be in equilibrium because of earlier demographic events, we allow for uncertainty in the initial conditions by initializing each iteration in the present model using a coalescent-based allele frequency distribution determined by a value (*θ*) randomly drawn from a uniform distribution. A population of size Ne is created and randomly mated for *t* ∈ [2, 8] generations, again with *t* drawn from a uniform distribution. Progeny from the adults in generation *t* is sampled, and summary statistics are then calculated from this sample.

The simulation model samples each time from a uniform flat prior distribution for Ne ∈ [4, 400] (except where specified as different), *θ* ∈ (4.8 *×* 10^−5^, 4.8 *×* 10^−3^), and duration in generations *t* ∈ [2, 8] at size Ne, to generate *J* = 2 *×* 10^4^ values of the summary statistics. This prior is reasonable, as Ne can fall in this range, even for some populations with thousands of individuals, because Ne is often much smaller than the total population size [15].

#### 2.1.3 Estimating Ne using linear regression

In order to estimate the Ne value for the input population, we first simulate *J* populations as described in Subsection 2.1.2 and calculate the summary statistics for each population. We will denote the statistics for the *j*-th population as *S*_*j*_ for *j* = 1, …, *J*. Hence, each *S*_*j*_ is a vector of five real numbers. Next, we denote the summary statistics for the input population as *S*^∗^. We note that *S*^∗^ is also a vector of five real numbers. Next, we compute the Euclidean distance between the *S*_*j*_ and *S*^∗^ for each *j* = 1, …, *J*. All datasets that have an Euclidean distance less than *δ*, which is denoted as a distance threshold used to regulate the number of acceptable candidates, are used to estimate the target Ne value. This estimation is accomplished using weighted local regression. We refer the reader to Tallmon et al. [30] for more details about this step.

Lastly, we use a Box-Cox transformation (*λ* = −0.2) of Ne in all regressions in order to ensure *θ*_*i*_, described in the previous subsection, are robust to changes in *δ*. Values of Ne accepted within 0.02 are then regarded as samples from the posterior distribution of Ne.

### 2.2 Implementation

We implemented ONeSAMP 3.0 in Python. All simulations were run on a server with AMD EPYC 7702 CPU, running the Red Hat Enterprise Linux 7.7 (64-bit, kernel 3.10.0). Our method takes as input an input dataset in GENEPOP format, a lower Ne value (`-lNe` parameter), an upper Ne value (`-uNe` parameter), a minimum allele frequency (`-m` parameter), a mutation rate value (`-r` parameter), lower and upper *θ* range (`-lt` and `-ut` parameters), lower and upper duration range (`-ld` and `-ud` parameters), the number of repeated simulation trails value (`-s` parameter), missing data rate for individual size and loci size (`-i` and `-l` parameters), and the input file name (`-o` parameter). The default values are as follows: minimal allele frequency: 0.05, mutation rate: 1.2 *×* 10^−8^, Ne range: {*x* ∈ ℤ | 50 ≤ *x* ≤ 150}, *θ* range which is the population mutation rate: {*x* ∈ ℤ | 4.8 *×* 10^−3^ ≤ *x* ≤ 4.8 *×* 10^−5^}, ONeSAMP 3.0 trial size: 2 *×* 10^4^, duration range: {*x* ∈ Z | 2 ≤ *x* ≤ 8}, and both the loci and the individual missing rate was set as 20%. Using a combination of real and simulated SNP data, we compare the performance of ONeSAMP 3.0 to that of `LDNe`.

#### 2.2.1 Data description

We used various datasets to measure the performance of ONeSAMP 3.0 and `LDNe`. First, we simulated population data with an increasing number of individuals and loci, and a target Ne value of 100. We generated populations with 50, 100, and 200 individuals and, for each population, we varied the number of loci to be 40, 80, 160, or 320. In addition, we simulated datasets with the same number of individuals and loci, with a target Ne value of 200. We used the default parameter settings to generate the initial simulated dataset but modified the Ne range to be {*x* ∈ ℤ | 50 ≤ *x* ≤ 150}, {*x* ∈ ℤ | 20 ≤ *x* ≤ 250} for Ne = 100, and {*x* ∈ ℤ | 150 ≤ *x* ≤ 250} for Ne = 200.

#### 2.2.2 Results of simulated data with Ne equal to 100

Both ONeSAMP 3.0 and `LDNe` were able to infer the value of Ne for 12 datasets. To minimize bias, we simulated each set of simulated conditions 30 times, each time computing the median predicted Ne value along with the upper and lower 95% quantile. These results are shown in Figure 1, which illustrates that ONeSAMP 3.0 consistently provided accurate predictions with narrow intervals for the upper and lower 95% quantile. We calculated the difference between the predicted Ne by ONeSAMP 3.0 and the target Ne value across all experiments with target Ne value of 100. When the number of individuals was 50, 100, and 200, the mean prediction of ONeSAMP 3.0 was 105.3, 105.9, and 106.6, with confidence intervals having sizes 74.2, 57.3, and 38.4, respectively. Hence, we witnessed that our method had increasingly narrow Ne confidence intervals as the number of individuals increases.

**Figure 1.**
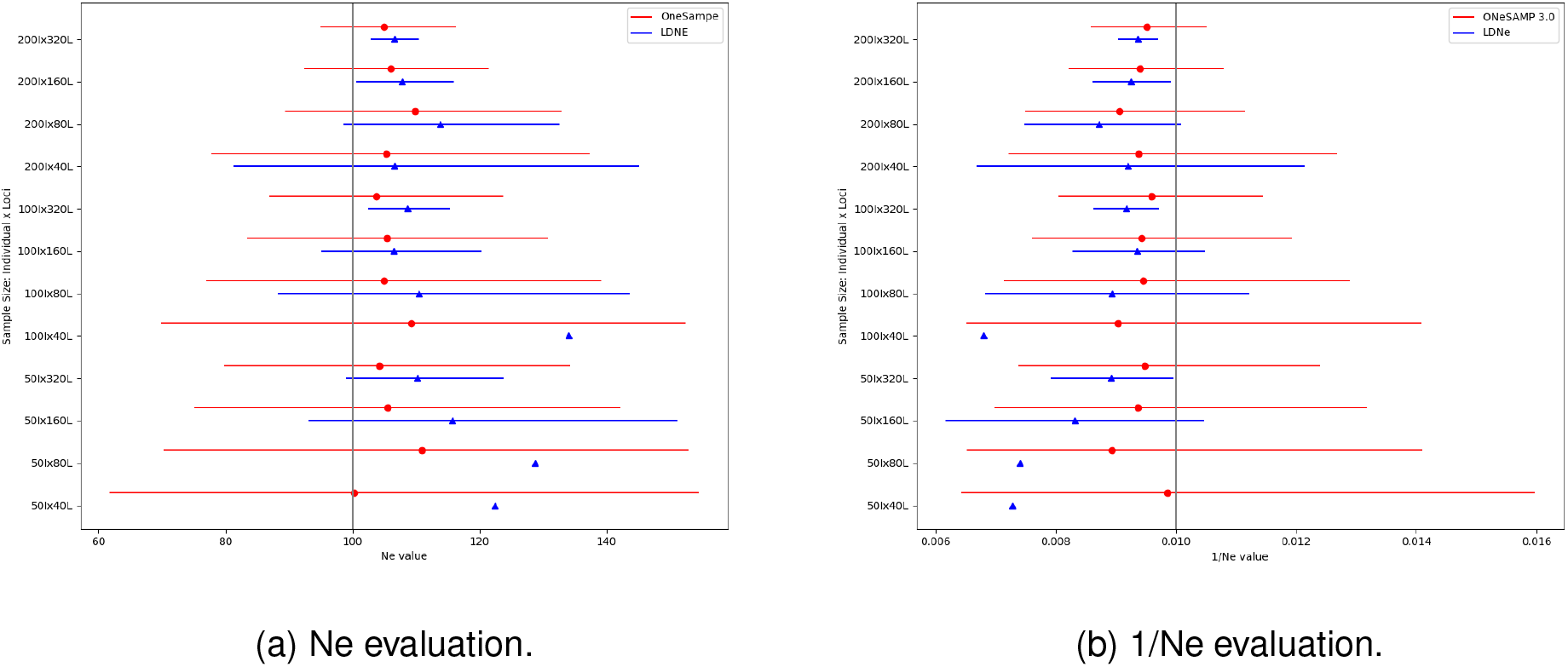
Estimator performance when Ne = 100, Ne range value with {*x* ∈ ℤ | 50 ≤ *x* ≤ 150}. Each point corresponds to the median Ne value, and the horizontal line illustrates the confidence interval. Two adjacent horizontal lines depict the Ne estimates for the same population by ONeSAMP 3.0 (red) and `LDNe` (blue). The absence of a line signifies an infinite confidence interval for that population.

Next, we studied the size of the confidence interval with an increasing number of loci and a larger population size. The confidence interval of ONeSAMP 3.0 narrowed considerably with a larger number of loci. When the number of individuals was 200, we observed the confidence interval had a steady decrease of 64.1% from 40 loci to 320 loci. Similarly, the confidence interval decreased from 82.5 to 36.8 when the number of loci went from 40 to 320 and the number of individuals was 100, and it decreased from 92.8 to 54.3 when the number of loci went from 40 to 320 and the number of individuals was 50. Thus, we see that the confidence intervals decreased when the number of individuals and/or the number of loci increased, which is reflective of the Law of Large Numbers [37], i.e., increasing the number of loci or individuals sampled will increase the convergence to the median Ne.

Lastly, we calculated the difference between the predicted Ne value and the target Ne value, which we denote this as *χ*, and determined how this parameter changes as the number of loci and individuals increases. We illustrate the trends in *χ* in Figure 3, which displays difference between the target Ne value and the predicted Ne value with increasing quantities of loci. We show that ONeSAMP 3.0 consistently produced small *χ* values across all scenarios and observed that increasing the number of loci leads to improved accuracy.

**Figure 2.**
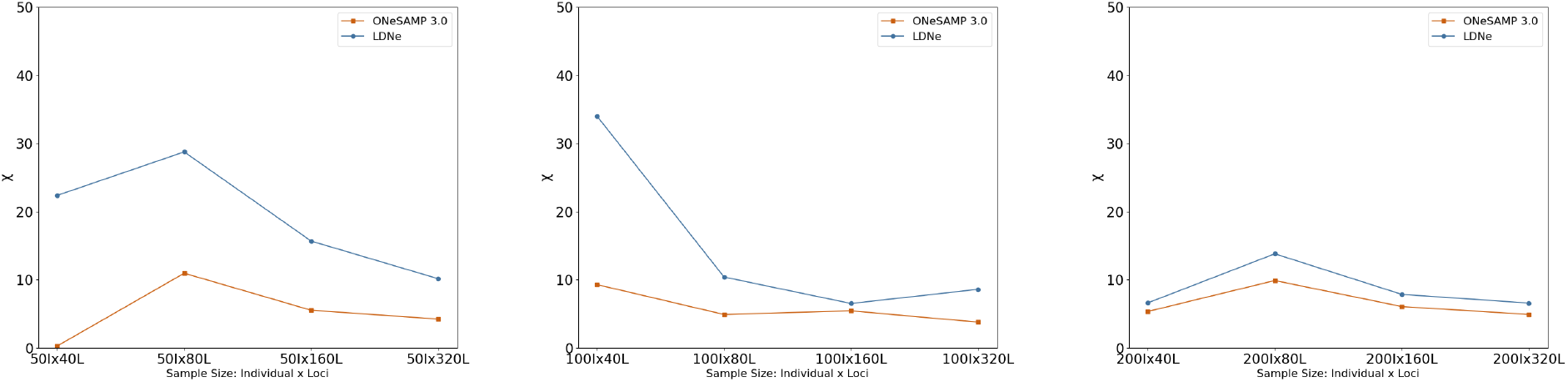
This diagram shows the sample mean difference between the target Ne value and the predicted Ne value. The difference in the mean is calculated by running both ONeSAMP 3.0 and `LDNe` 30 times on 12 different simulated datasets, and calculating the mean. The target Ne is 100.

**Figure 3.**
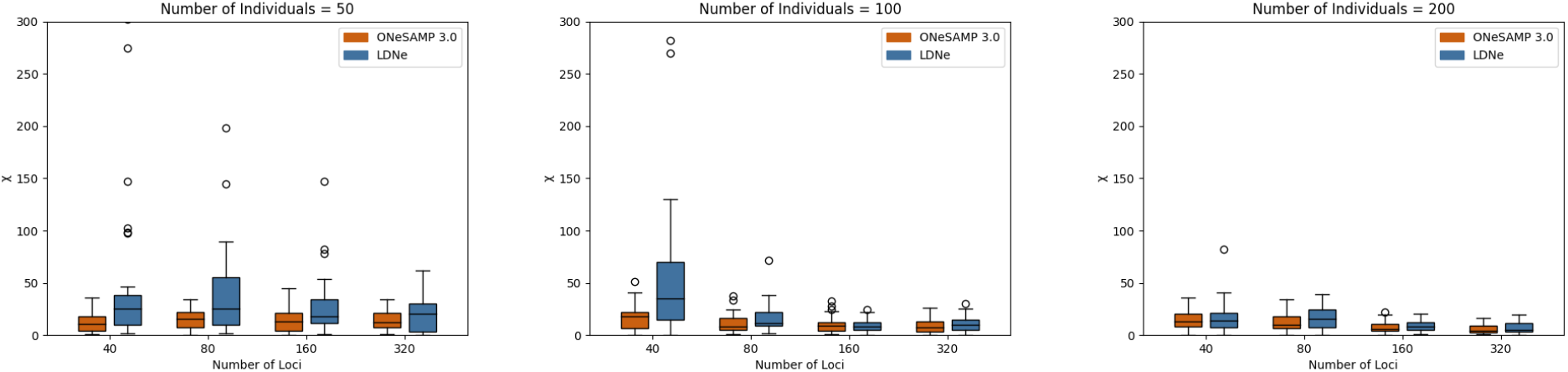
This diagram displays the difference between the target Ne value and the predicted Ne value with increasing quantities of loci (i.e., the number of loci is 40, 80, 160, and 320) and various population sizes. The graphs from left to right illustrate the difference with 50, 100, and 200 individuals, respectively.

#### 2.2.3 Results of simulated data with Ne equal to 200

We conducted a performance evaluation of ONeSAMP 3.0 using populations that have a higher target Ne value. Figure 4 illustrates that as the number of individuals and loci increases, both ONeSAMP 3.0 and `LDNe` produce more precise predictions. ONeSAMP 3.0 performs well in scenarios with limited data. For example, with 50 individuals, and fewer than 320 loci, and with 100 individuals and fewer than 40 loci, ONeSAMP 3.0 achieved results as accurate as 0.065% and 2.735% above the target Ne value, respectively. As previously seen with a target Ne value of 100, the confidence interval of ONeSAMP 3.0 decreases as the number of loci increases.

**Figure 4.**
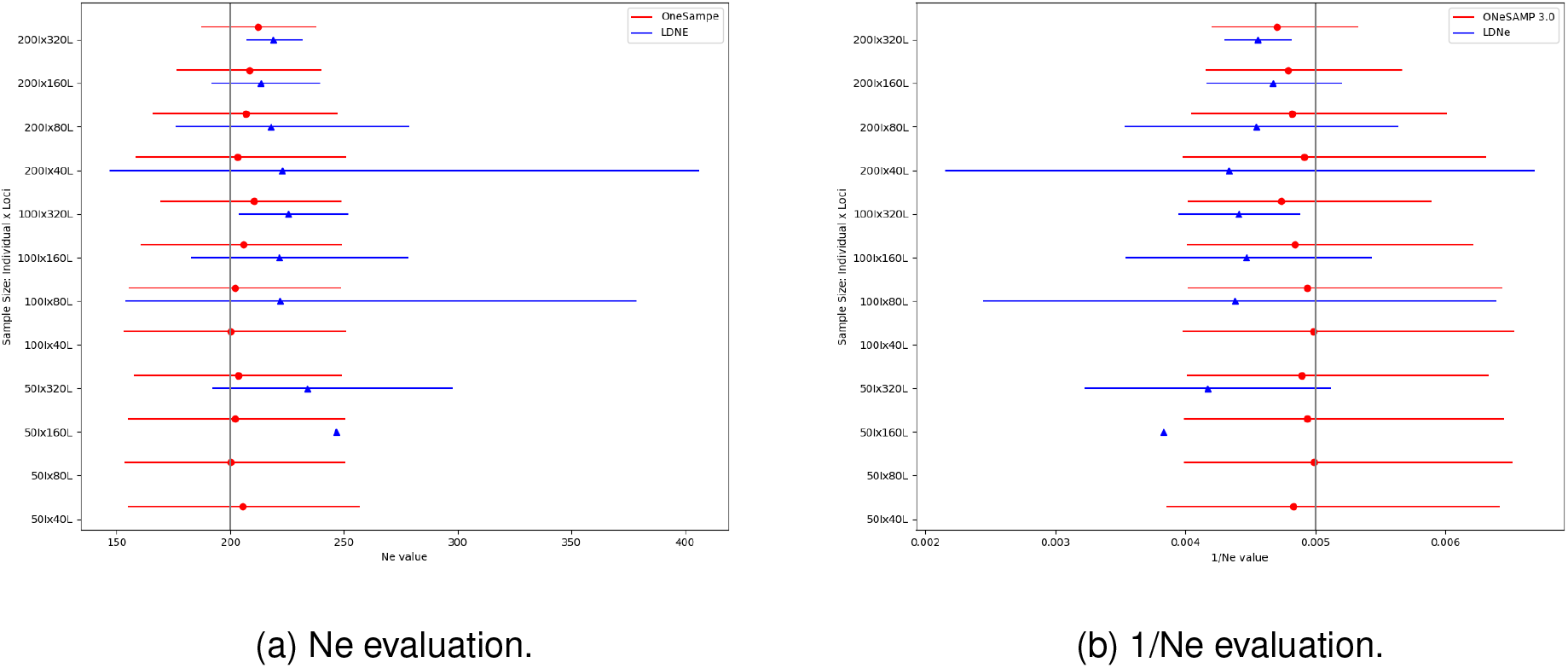
Results from simulations with all populations having a target Ne value of 200, Ne range equal to {*x* ∈ ℤ | 150 ≤ *x* ≤ 250}. Each point corresponds to the median Ne value, and the horizontal line illustrates the confidence interval. Two adjacent horizontal lines depict the Ne estimates for the same population by ONeSAMP 3.0 (red) and `LDNe` (blue). If a line or point is absent, it indicates that the confidence interval or predicted value of Ne for that population is infinite.

Figure 5 shows that ONeSAMP 3.0 consistently produces small and stable *χ* values, indicating its reliability in making predictions. Furthermore, we observed that increasing the number of loci results in improved accuracy, as illustrated in Figure 6. In addition, this figure shows that ONeSAMP 3.0 yields consistently narrow confidence intervals and few outlier values.

**Figure 5.**
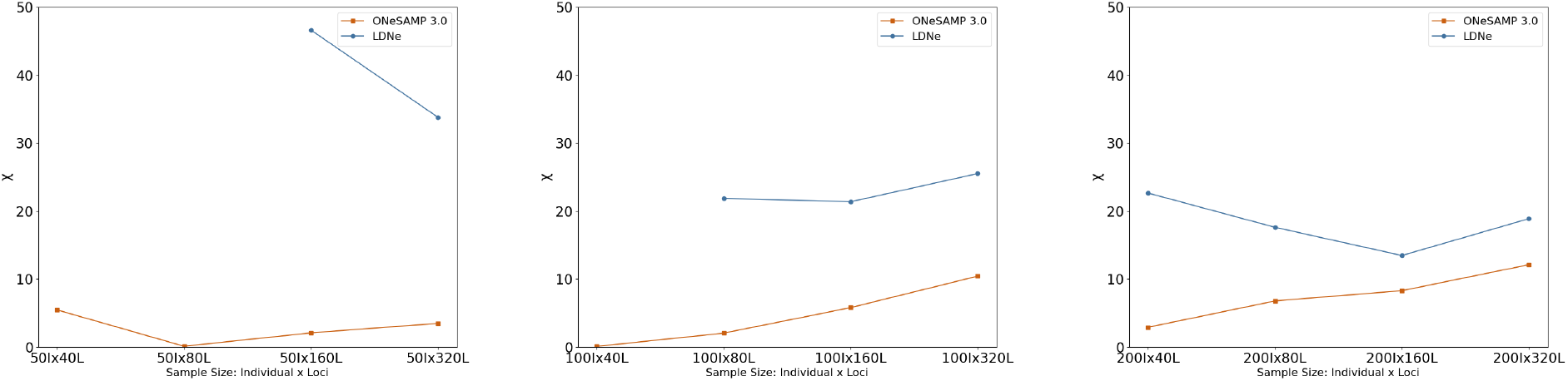
This diagram shows the mean difference between the target Ne value and the predicted Ne value. The mean difference is calculated by running either ONeSAMP 3.0 and `LDNe` 30 times on 12 different simulated datasets and calculating the mean. The target Ne is 200.

**Figure 6.**
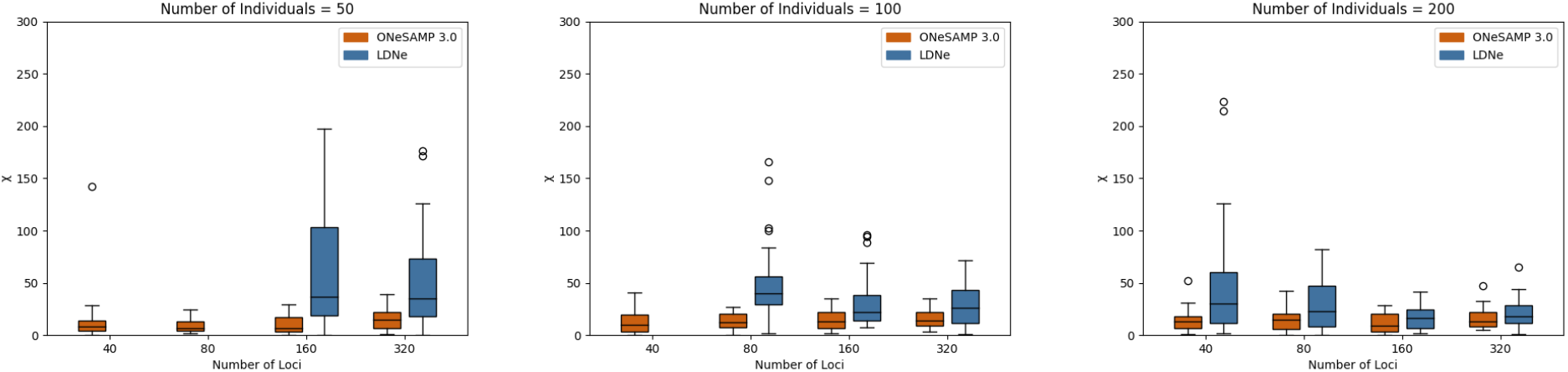
This diagram displays how the difference between the target Ne value and the predicted Ne value with increasing loci (i.e., the number of loci is 40, 80, 160, and 320) and various population sizes. The leftmost graph illustrates the difference with 50 individuals. The middle graph illustrates the difference with 100 individuals. And finally, the rightmost graph illustrates the difference with 200 individuals. The target Ne is 200.

#### 2.2.4 Results of simulated data with a narrow Ne range

Based on the preceding experimental outcomes, we see that ONeSAMP 3.0 demonstrates consistent favorable performance across a variety of datasets. Motivated by these findings, we conducted an additional experimental inquiry aimed at elucidating the performance dynamics of ONeSAMP 3.0 under an extended scope of Ne values, spanning from 20 to 250. As illustrated in Figure 7, it is discerned that in scenarios involving modestly sized datasets, a distinct trend emerges: Within the experimental context—specifically when the number of individuals is less than 100 and the loci size is under 80—ONeSAMP 3.0 outperforms the `LDNe` but `LDNe` outperforms ONeSAMP 3.0 on all other datasets. Hence, the results show that `LDNe` and ONeSAMP 3.0 produce improved predictions but in different scenarios. Moreover, we see that the mean of the predicted Ne values of both ONeSAMP 3.0 and `LDNe` is comparable; with only negligible difference between the values. More dramatic differences are shown between the confidence intervals with ONeSAMP 3.0 having considerably larger confidence intervals than `LDNe` which grow narrower as the number of individuals or loci increases. There was only a single dataset—our smallest dataset with 50 individuals and 40 loci—where both methods struggled to perform accurate and consistent results.

**Figure 7.**
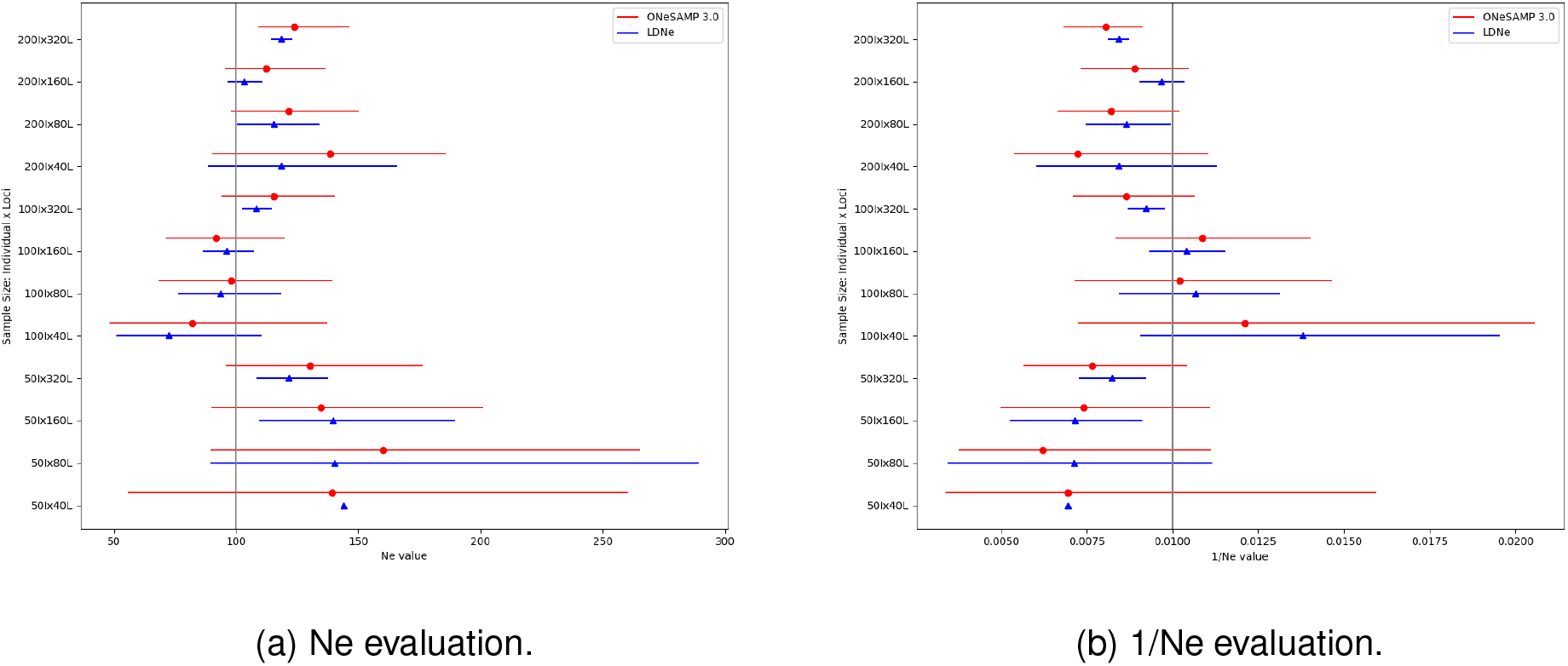
Estimator performance when Ne = 100, Ne range value with {*x* ∈ ℤ | 20 ≤ *x* ≤ 250}. Each point corresponds to the median Ne value, and the horizontal line illustrates the confidence interval. Two adjacent horizontal lines depict the Ne estimates for the same population by ONeSAMP 3.0 (red) and `LDNe` (blue). The absence of a line signifies an infinite confidence interval for that population.

#### 2.2.5 Empirical dataset

To provide an example of the use and interpretation of ONeSAMP 3.0 as a genetic estimator of Ne, we reanalysed data from six sub-populations of Channel Island foxes sampled across their range including San Miguel Island (SMI), Santa Rosa Island (SRI), Santa Cruz Island (SCI), San Nicolas Island (SNI), Santa Catalina Island (SCA), and San Clemente Island (SCL) at 4,860 SNPs [38]. This data set was chosen because several of the Channel Island fox sub-populations experienced population bottlenecks with varying levels of severity in the late 1990’s due to heavy predation by invasive golden eagles [39, 40] and disease [41]. In this experiment, we first filtered all loci containing missing data. The number of loci that remained after filtering is shown in Table 1. SNI sub-population did not retain sufficient enough loci were not included in our analyses. Next, we further reduced the number of SNPs by randomly sampling 50, 100, 200, and 300 loci without replacement to compare the performance of ONeSAMP 3.0 and `LDNe` with limited loci.

**Table 1:**
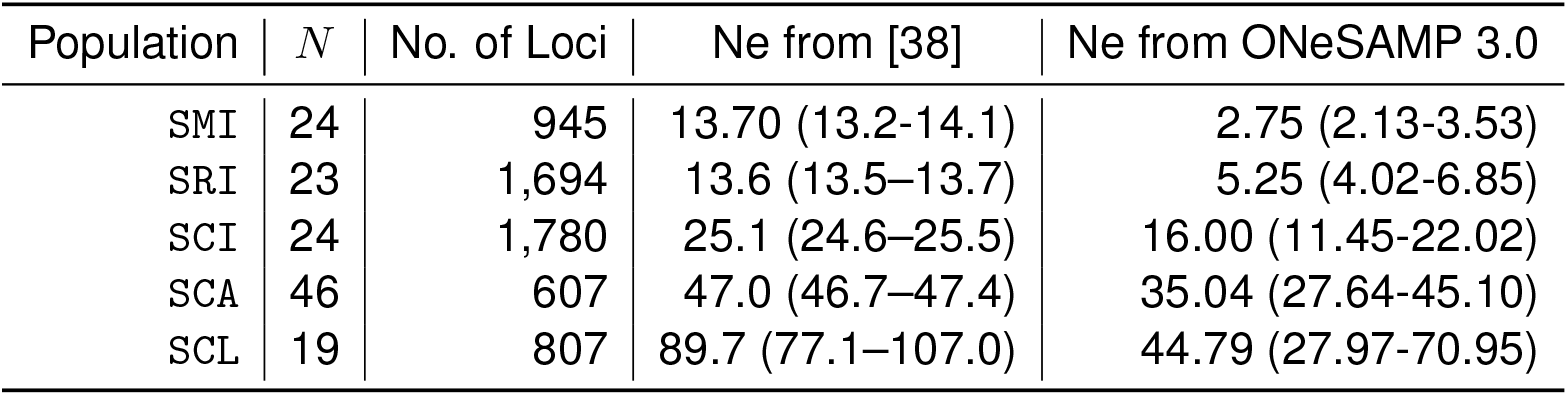
A table giving the number of individuals in each dataset (*N*), the number of loci after filtering, the value of Ne predicted by [38], and the value of Ne predicted by ONeSAMP 3.0. There were 4,860 loci in the original dataset. The confidence intervals are given along with the predicted Ne values. The number of loci used for these predictions was 300.

| Population | $N$ | No. of Loci | Ne from [38] | Ne from ONeSAMP 3.0 |
| --- | --- | --- | --- | --- |
| SMI | 24 | 945 | 13.70 (13.2-14.1) | 2.75 (2.13-3.53) |
| SRI | 23 | 1,694 | 13.6 (13.5-13.7) | 5.25 (4.02-6.85) |
| SCI | 24 | 1,780 | 25.1 (24.6-25.5) | 16.00 (11.45-22.02) |
| SCA | 46 | 607 | 47.0 (46.7-47.4) | 35.04 (27.64-45.10) |
| SCL | 19 | 807 | 89.7 (77.1-107.0) | 44.79 (27.97-70.95) |

#### 2.2.6 Results of empirical data

Our results show that when Ne is small, ONeSAMP 3.0 produces estimates of Ne that are comparable to `LDNe`; See Figure 8 and Table 1. Conversely, when Ne is large, ONeSAMP 3.0 produces small confidence intervals when the number of loci is less than 200. While both ONeSAMP 3.0 and `LDNe` produced infinite estimates for Ne for SMI, SCL, and SCI populations with 50 loci, our method provided more accurate results regularly when the number of loci and individuals sampled was small.

**Figure 8.**
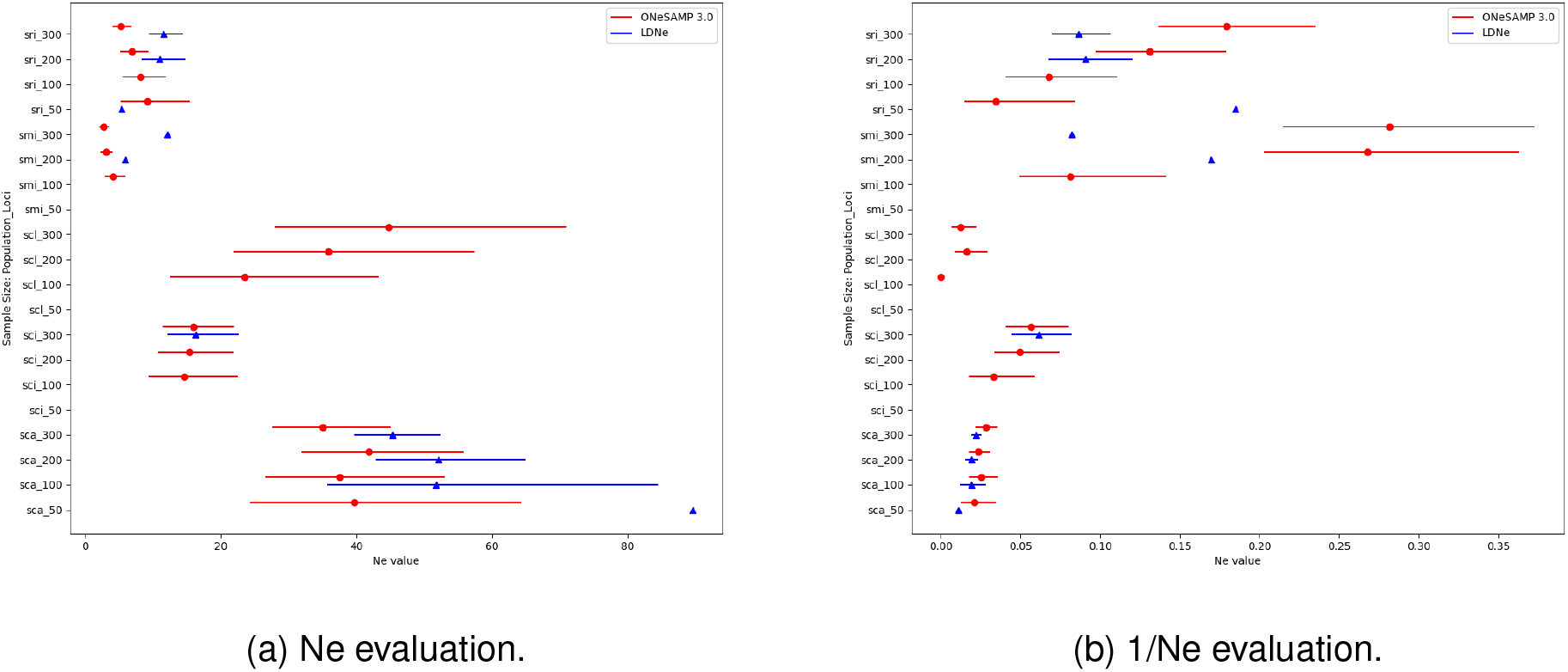
This diagram depicts the outcomes of our experiment with 35 populations. Each horizontal line shows the estimated range of Ne for a particular population, and each pointer reflects the estimated median Ne for that population. Two adjacent horizontal lines depict the Ne estimates for the same population by ONeSAMP 3.0 (red) and `LDNe` (blue).

## 3 Discussion

We present ONeSAMP 3.0, which is a novel approximate Bayesian computation method that predicts accurate estimates of Ne from a single population sample. Through simulations and empirical data, we demonstrate ONeSAMP 3.0 is optimized to estimate Ne reliably even when there exists a limited number of loci and individuals. Furthermore, ONeSAMP 3.0 is flexible enough to obtain estimates of Ne that are robust to population demographic history and sampling designs. More specifically, we showed that the accuracy of ONeSAMP 3.0’s has less than a 6% bias when Ne is small, i.e., equal to 100, and less than a 5% bias when Ne is larger, i.e., when Ne is equal to 200. Moreover, we showed that increasing the number of loci resulted in a consistent decrease in the width of the confidence interval. However, our experimental results also indicate that, in cases where the dataset is small (i.e., the number of individuals is below 100 and the number of loci is less than 160), the extended Ne range consistently leads to improved performance of the ONeSAMP 3.0 model. In optimal circumstances, the ONeSAMP 3.0 model demonstrates a significant enhancement, with augmentation rates between 35% and 66%.

Our results using empirical data from island fox sub-populations show that both ONeSAMP 3.0 and `LDNe` produced infinite estimates Ne for SMI, SCL, and SCI populations with 50 loci. This is likely due to a lack of sufficient genetic variation in these sub-populations to produce a reliable estimate with only 50 loci [38]. However, ONeSAMP 3.0 does produce accurate results even in cases when there are a limited number of polymorphic loci and samples available; see Figure 8. This accuracy can be partially attributed to the generation of a large number of simulated populations, from which the probability distributions and related summary statistics of Ne under different scenarios are calculated. This Bayesian approach allows ONeSAMP 3.0 to integrate over the uncertainty in the parameters (i.e., target Ne values, values for *θ*, and mutation rate) to obtain a robust posterior distribution for Ne. This is in contrast to competing methods (such as `LDNe`), which only uses information from SNP data and/or linkage information [24]. Information on the size or history of a population can also be easily incorporated into ONeSAMP 3.0’s prior distribution for Ne. Moreover, because the prior distribution is generated by sampling simulated populations with the same number of loci and individuals as the empirical data set, ONeSAMP 3.0 should mitigate the detrimental effects of small sample size on estimates of Ne. In addition, we note that one important feature of a Bayesian approach is that it limits the posterior confidence interval by use of a prior, which is required as input. Arguably, non-Bayesian approaches (such as `LDNe`) could truncate the confidence intervals down to known or finite upper limits in the cases where estimates are infinite. The results reported here are directly from `LDNe` and ONeSAMP 3.0 when provided identical simulated and real datasets.

Next, we note that when new genotype or allele frequency summary statistics become available, it would be simple to adapt the approximate Bayesian method outlined here to estimate Ne. As an increasing number of molecular markers become available for a particular taxon, the approximate Bayesian framework provides an attractive and efficient method for estimating parameter values from the existing data. Nonetheless, it appears that reliable and exact Ne estimate is attainable using ONeSAMP 3.0 if moderate numbers of individuals and independent loci are sampled. Further research may also find more informative summary statistics to optimize this technique for application with various molecular marker types.

In conservation genetic practices, estimates of Ne provide researchers with important information for monitoring and recovery planning of populations [12, 42]. The general rule of thumb that an Ne of less than 50 should be considered a short-term extinction risk and an Ne above 500 poses no immediate genetic threat of extinction has garnered considerable support [8, 43]. However, a general problem for estimating Ne with genetic data is that each method has underlying assumptions regarding the sampled population’s demography and underlying population structure [7, 17]. Therefore, each estimate of Ne must be interpreted with caution in cases of at-risk species regardless of the methodology. Lastly, we conclude by pointing out that it is important to correct Ne estimates for the number of chromosomes of the given study species when you estimate Ne for real populations. This can be accomplished by using the corrections in Waples et al. [44]. Another option is to preprocess the data so that the method only uses pairs of loci on different chromosomes.

## 4 Recommendations

Our research findings suggest that the prediction of ONeSAMP 3.0 performs well when Ne value is large relative to the size of the dataset, i.e., as tested at Ne = 100 and Ne = 200. This becomes particularly pronounced when the dataset consists of a limited number of individuals (as tested with 50 individuals), a scenario where other methods may fail to produce reasonable estimates. On the other hand, the ONeSAMP 3.0 predictor can generate precise predictions in these instances. We note that ONeSAMP 3.0 can efficiently make predictions for a small number of SNPs—i.e., 500 or less loci—but slows down considerably as the number of SNPs surpasses 500. For this reason, we recommend that LDNe be used when the dataset has a large number of loci.

## 5 Conclusion

Generating estimates of Ne from genetic data continues to be a highly cost-effective and reliable approach. Our method only requires a standard GENEPOP file and should be accessible to researchers familiar with basic Linux or UNIX scripting. We have demonstrated that ONeSAMP 3.0 can provide a more accurate prediction than `LDNe` when the number of individuals and/or SNPs is relatively small. Moreover, ONeSAMP 3.0 is able to provide accurate estimates of Ne even for small datasets, which will be useful to researchers interested in exploring Ne with limited genetic resources or sample sizes available. We anticipate ONeSAMP 3.0 will become an important exploratory tool for estimating Ne in non-model species. The development of ONeSAMP 3.0 also opens the door for future considerations including the addition of statistics that main increase the accuracy of the prediction, and parallelization of the computation that will increase the efficiency of the tool. Lastly, the implementation also enables other machine learning models to used in replace of linear regression. In particular, using a non-linear machine learning model—such as a random forest classifier or convolutional neural network—might lead to some notable performance gains and increased precision.

## Acknowledgements

We thank Dr. Robin Waples for his helpful suggestions and discussions of Ne estimation and feedback on an earlier draft of this manuscript.

## References

[1] J.F. Crow. Mathematical Topics in Population Genetics, Volume 1. 1970.

[2] R. Lande, S. Engen, and B.-E. Saether. Optimal harvesting of fluctuating populations with a risk of extinction. The American Naturalist, 145(5):728–745.

[3] M. Lynch, J. Conery, and R. Burger. Mutation accumulation and the extinction of small populations. The American Naturalist, 146(4):489–518, 1995.

[4] R.S. Waples. Life-history traits and effective population size in species with overlapping generations revisited: the importance of adult mortality. Heredity, 117(4):241–250, 2016.

[5] R.S. Waples. Making sense of genetic estimates of effective population size. Molecular Ecology, 25(19):4689–4691, 2016.

[6] R.S. Waples. What is Ne, anyway? Journal of Heredity, 113(4):371–379, 2022.

[7] J. Wang, E. Santiago, and A. Caballero. Prediction and estimation of effective population size. Heredity, 117(4):193–206, 2016.

[8] M.E. Soule and B.A. Wilcox. Conservation Biology. An Evolutionary-Ecological Perspective. Sinauer Associates, Inc., 1980.

[9] R. Frankham, C.J.A. Bradshaw, and B.W. Brook. Genetics in conservation management: revised recommendations for the 50/500 rules, red list criteria and population viability analyses. Biological Conservation, 170:56–63, 2014.

[10] J.T. Howard, J.E. Pryce, C. Baes, and C. Maltecca. Inbreeding in the genomics era: Inbreeding, inbreeding depression, and management of genomic variability. Journal of Dairy Science, 100(8):6009–6024, 2017.

[11] H. Ellegren and N. Galtier. Determinants of genetic diversity. Nature Reviews Genetics, 17(7):422–433, 2016.

[12] S. Hoban, M. Bruford, J. Jackson D’Urban, M. Lopes-Fernandes, M. Heuertz, P.A. Hohenlohe Paz-Vinas, P. Sjö gren-Gulve, G. Segelbacher, C. Vernesi, et al. Genetic diversity targets and indicators in the CBD post-2020 Global Biodiversity Framework must be improved. Biological Conservation, 248:108654, 2020.

[13] R. Bijlsma, J. Bundgaard, and A.C. Boerema. Does inbreeding affect the extinction risk of small populations?: predictions from Drosophila. Journal of Evolutionary Biology, 13(3):502–514, 2000.

[14] D. Newman and D. Pilson. Increased probability of extinction due to decreased genetic effective population size: experimental populations of clarkia pulchella. Evolution, 51(2):354–362, 1997.

[15] F.P. Palstra. Determinants and Evolutionary Consequences of Effective Population Size in Atlantic Salmon (Salmo Salar). Dalhousie University, 2008.

[16] F.P. Palstra and D.J. Fraser. Effective/census population size ratio estimation: a compendium and appraisal. Ecology and Evolution, 2(9):2357–2365, 2012.

[17] R.S. Waples. Genetic estimates of contemporary effective population size: to what time periods do the estimates apply? Molecular Ecology, 14(11):3335–3352, 2005.

[18] T. Nomura. Estimation of effective number of breeders from molecular coancestry of single cohort sample. Evolutionary Applications, 1(3):462–474, 2008.

[19] O.L. Zhdanova and A.I. Pudovkin. Nb hetex: a program to estimate the effective number of breeders. Journal of Heredity, 99(6):694–695, 2008.

[20] J. Wang. A new method for estimating effective population sizes from a single sample of multilocus genotypes. Molecular Ecology, 18(10):2148–2164, 2009.

[21] R.S. Waples and R.K. Waples. Inbreeding effective population size and parentage analysis without parents. Molecular Ecology Resources, 11:162–171, 2011.

[22] M. Barbato, P. Orozco-terWengel, M. Tapio, and M.W. Bruford. SNeP: a tool to estimate trends in recent effective population size trajectories using genome-wide SNP data. Frontiers in Genetics, 6:109, 2015.

[23] Waples, R. S., and C. Do. LDNe: a program for estimating effective population size from data on linkage disequilibrium. Molecular Ecology Resources, 2008.

[24] C. Do Waples, R. S., Peel, D. G. M. Macbeth, B. J. Tillett, and J. R. Ovenden. NeEstimator V2: re-implementation of software for the estimation of contemporary effective population size (Ne) from genetic data. Molecular Ecology Resources, 2014.

[25] S. Tavaré, D.J. Balding, R.C. Griffiths, and P. Donnelly. Inferring coalescence times from DNA sequence data. Genetics, 145(2):505–518, 1997.

[26] J.K. Pritchard and N.A. Rosenberg. Use of unlinked genetic markers to detect population stratification in association studies. The American Journal of Human Genetics, 65(1):220–228, 1999.

[27] S.A. Tishkoff, R. Varkonyi, N. Cahinhinan, S. Abbes, G. Argyropoulos, G. Destro-Bisol, A. Drousiotou, B. Dangerfield, G. Lefranc, J. Loiselet, A. Piro, M. Stoneking, A. Tagarelli, G. Tagarelli, E.H. Touma, S.M. Williams, and A.G. Clark. Haplotype diversity and linkage disequilibrium at human G6PD: recent origin of alleles that confer malarial resistance. Science, 293(5529):455–462, 2001.

[28] N. Nikolic and C. Chevalet. Detecting past changes of effective population size. Evolutionary Applications, 7(6):663–681, 2014.

[29] M.A. Beaumont, W. Zhang, and D.J. Balding. Approximate bayesian computation in population genetics. Genetics, 162(4):2025–2035, 2002.

[30] D.A. Tallmon, A. Koyuk, G. Luikart, and M.A. Beaumont. COMPUTER PROGRAMS: ONe-SAMP: a program to estimate effective population size using approximate Bayesian computation. Molecular Ecology Resources, 8(2):299–301, 2008.

[31] M. Nei. Molecular Evolutionary Genetics. Columbia University Press, 1987.

[32] AHD Brown, MW Feldman, and E Nevo. Multilocus structure of natural populations of hordeum spontaneum. Genetics, 96(2):523–536, 1980.

[33] B.S. Weir. Inferences about linkage disequilibrium. Biometrics, 1:235–254, 1979.

[34] W.G Hill. Estimation of effective population size from data on linkage disequilibrium. Genetics Research, 38(3):209–216, 1981.

[35] D.A. Tallmon, G. Luikart, and M.A. Beaumont. Comparative evaluation of a new effective population size estimator based on approximate bayesian computation. Genetics, 167(2):977–988, 2004.

[36] F. Balloux. Heterozygote excess in small populations and the heterozygote-excess effective population size. Evolution, 58(9):1891–1900, 2004.

[37] Michel Dekking. A Modern Introduction to Probability and Statistics. Springer, 2005.

[38] W.C. Funk, R.E. Lovich, P.A. Hohenlohe, C.A. Hofman, S.A. Morrison, S.T. Sillett, C.K. Ghalambor, J.E. Maldonado, T.C. Rick, M.D. Day, N.R. Polato, S.W. Fitzpatrick, T.J. Coonan, K.R. Crooks, A. Dillon, D.K. Garcelon, J.L. King, C.L. Boser, N. Gould, and W.F. Andelt. Adaptive divergence despite strong genetic drift: genomic analysis of the evolutionary mechanisms causing genetic differentiation in the island fox (Urocyon littoralis). Molecular Ecology, 25(10):2176–2194, 2016.

[39] G.W. Roemer, T.J. Coonan, D.K. Garcelon, J. Bascompte, and L. Laughrin. Feral pigs facilitate hyperpredation by golden eagles and indirectly cause the decline of the island fox. Animal Conservation, 4(4):307–318, 2001.

[40] T.J. Coonan, C.A. Schwemm, and D.K. Garcelon. Decline and recovery of the island fox: a case study for population recovery. Cambridge University Press, 2010.

[41] S.F. Timm, L. Munson, B.A. Summers, K.A. Terio, E.J. Dubovi, C.E. Rupprecht, S. Kapil, and D.K. Garcelon. A suspected canine distemper epidemic as the cause of a catastrophic decline in Santa Catalina Island foxes (Urocyon littoralis catalinae). Journal of Wildlife Diseases, 45(2):333–343, 2009.

[42] G. Luikart, N. Ryman, D.A. Tallmon, M.K. Schwartz, and F.W. Allendorf. Estimation of census and effective population sizes: the increasing usefulness of DNA-based approaches. Conservation Genetics, 11:355–373, 2010.

[43] I.G. Jamieson and F.W. Allendorf. How does the 50/500 rule apply to MVPs? Trends in Ecology & Evolution, 27(10):578–584, 2012.

[44] Ryan K Waples, Wesley A Larson, and Robin S Waples. Estimating contemporary effective population size in non-model species using linkage disequilibrium across thousands of loci. Heredity, 117(4):233–240, 2016.

